# A yeast model of 5-oxoproline accumulation reveals a general toleration to 5-oxoproline

**DOI:** 10.64898/2025.12.21.695865

**Authors:** Pratiksha Dubey, B S Vaishnavi, Praveen Singh, Shantanu Sengupta, Anand K Bachhawat

## Abstract

5-oxoproline (5-OP) or pyroglutamic acid is an intermediate of the degradation arc of the glutathione cycle. It is metabolized into glutamate through the action of the 5-oxoprolinase enzyme, the only enzyme known to act on this metabolite. 5-OP has long been known to be relatively inert with a proposed role as an osomoprotectant. Recent studies on the 5-oxoprolinase enzyme in mammalian cells have, however, shown that knockdown or deletions of the 5-oxoprolinase make mice (and humans) prone to heart failure. This was ascribed to oxidative stress due to a two-fold elevation in 5-OP. To examine the consequences of 5-oxoproline accumulation more rigorously, we have created a yeast model for 5-oxoproline accumulation. We observed retardation of growth only when intracellular levels of 5-OP were increased 12-20 fold over the normal levels. A transcriptomics study was carried out under these conditions. We observed that while a large number of genes were regulated upto 2-fold, there were no single prominent pathways amongst them. Among the key genes upregulated were different efflux pumps. Using knockouts and also overexpression of selected genes, we could observe that many of the upregulated genes were involved in the cellular response to 5-OP accumulation. However, it did not appear that there was any significant oxidative stress response. The study suggests a need to reevaluate previous suppositions of the 5-OP induced oxidative stress response. Instead, we propose an alternative model to explain the possible consequences of 5-oxoprolinase deficiency based on these findings made with the yeast model.

## Introduction

5-oxoproline (5-OP), also known as pyroglutamic acid is widespread in nature (Kumar and Bachhawat, 2012). It is an intermediate of the glutathione cycle, a cycle which describes glutathione metabolism in living cells (Bachhawat and Yadav, 2018). 5-oxoproline can also be found at the N-terminus of some proteins which is formed from the spontaneous cyclization of N-terminal glutamate or glutamine in proteins (Abraham and Podell, 1981; Schilling et al., 2004). At the N-terminus sites of proteins 5-oxoproline contributes to both function and stability (Cummins and O’Connor, 1998). However, as a free metabolite its functions are less clear. It appears to have a role in glutamate storage and it has also been suggested to function as an osmoprotectant in microbes (Khmelenina et al., 1999; Kumar and Bachhawat, 2012; Trotsenko and Khelenina, 2002) and a moisturiser in mammals (Bouwstra et al., 2008; Wollenberg et al., 2025).

The only enzyme that acts on 5-oxoproline is 5-oxoprolinase. This enzyme hydrolyses 5-OP to yield glutamate. The enzyme is considered to be one of the most sluggish enzymes (Watanabe et.al., 2004). Thus 5-oxoproline in cells often accumulates and some of it is found to be secreted into the plasma (Van Der Werf et al., 1974) . Recent studies with both humans and mice have revealed that cardiac injury leads to 5-oxoprolinase depletion, resulting in elevated 5-OP levels in the plasma (Van Der Pol et al., 2018, 2017). The study also found that cardiac function after ischemic injury could be rescued by overexpressing 5-oxoprolinase (Van Der Pol et al., 2018, 2017). The study concluded that 5-OP accumulation was causing oxidative stress. This was surprising for two reasons. Firstly, 5-OP appears to be a relatively inert molecule, and its previous role has in fact been suggested in osmoprotection. Secondly, the increased levels in plasma observed during 5-oxoprolinase knockdown was only two-fold (Van Der Pol et al., 2018, 2017). This was not a very high increase in plasma level 5-OP since 5-OP levels are normally observed in plasma and levels are also known to vary with diet (Jackson et al., 1996).

It was important to understand therefore, if it was 5-oxoproline accumulation *per se* that might be important, or some other underlying mechanism (such as decreased intracellular glutathione). We have accordingly begun to investigate more carefully, whether 5-OP accumulation in cells does indeed cause oxidative stress. To address this question, we have created a yeast model for 5-OP accumulation. We initially tried creating a model where 5-OP would be generated endogenously. However, it could not lead to significant elevations in 5-OP. We finally established a model where a yeast strain lacking 5-oxoprolinase (*oxp1Δ* strain) was treated with excess 5-OP in the medium. Due to deletion of the 5-oxoprolinase gene, 5-OP cannot be metabolized leading to 5-OP accumulation. After establishing a model that could cause between 12- to 20- fold increase in 5-OP levels, the consequences of 5-OP accumulation were investigated using transcriptomics. RNA seq analysis followed by GO analysis revealed that xenobiotic export and efflux pathways were upregulated. In addition, transcription factors in diverse pathway were mildly upregulated including the stress response factor SKN7. In addition, several different enzymes and membrane proteins were upregulated. We confirmed using deletion strain that the knockout of many of these different genes, though not all, did in fact cause them to be more sensitive to 5-OP. This data suggests that 5-OP accumulation to 12-fold higher levels causes only a mild stress to the cell through multiple pathways which cumulatively leads to the growth defect in *oxp1Δ* strain.

## Results

### Creation of a yeast model for 5-oxoproline accumulation

To study the effect of intracellular accumulation of 5-OP in yeast cells we needed a system that could result in accumulation of a high amount of intracellular 5-OP. We initially conceived of two broad approaches. The first approach was to generate high levels of 5-OP endogenously by transforming into yeast a 5-OP generating enzyme, and the second approach was to exogenously provide different levels of 5-OP in the medium enabling its accumulations within the cell.

In the first approach we used heterologously expressed human Chac1. Chac1 degrades glutathione efficiently to generate 5-OP and Cys-gly (Kumar et al., 2012). We introduced this into a strain that was deleted for the endogenous glutathione degradation pathways Ecm38p and Dug3p. Ecm38p is a γ-glutamyl transpeptidase that cleaves glutathione to yield glutamate and Cys-gly and Dug3p is a component of the Dug2p–Dug3p complex that also yields glutamate and Cys-gly on glutathione degradation (Kaur et al., 2012). They were both distinct in their activity compared to Chac1 which cleaves glutathione to yield 5-OP and Cys-gly. Therefore, we used a strain background that is lacking both these enzymes. We also used the *met15Δ* background. This is an organic sulfur auxotroph and it enabled us to evaluate whether Chac1 was indeed degrading glutathione in the cells. Since the degraded glutathione would release cysteine to provide the sulfur source required. We also created a 5-oxoprolinase deletion (*oxp1Δ*) in this background. This stain, *met15Δdug3-2ecm38Δoxp1Δ* strain was transformed with either the *OXP1* expressing clone (TEF-*OXP1*) or with an empty vector. Those with TEF-*OXP1* clone would degrade 5-OP generated from Chac1, while those transformed with empty vector were expected to have 5-OP accumulated. We found that the vector transformed strains were showing slightly slower growth as compared to OXP1 overexpressed strains (Fig 1A). 5-OP levels were then estimated in the cell extracts using LC-MS. However, we could not observe any accumulation of 5-OP in the vector transformed cells (Fig 1B) Compared to the OXP1 transformed cells. In contrast there was some small increase in 5-OP levels in the OXP1 transformed cells in contrast to the expectations. This suggests that this model was not really suitable as a model for 5-OP accumulation.

**Figure 1.**
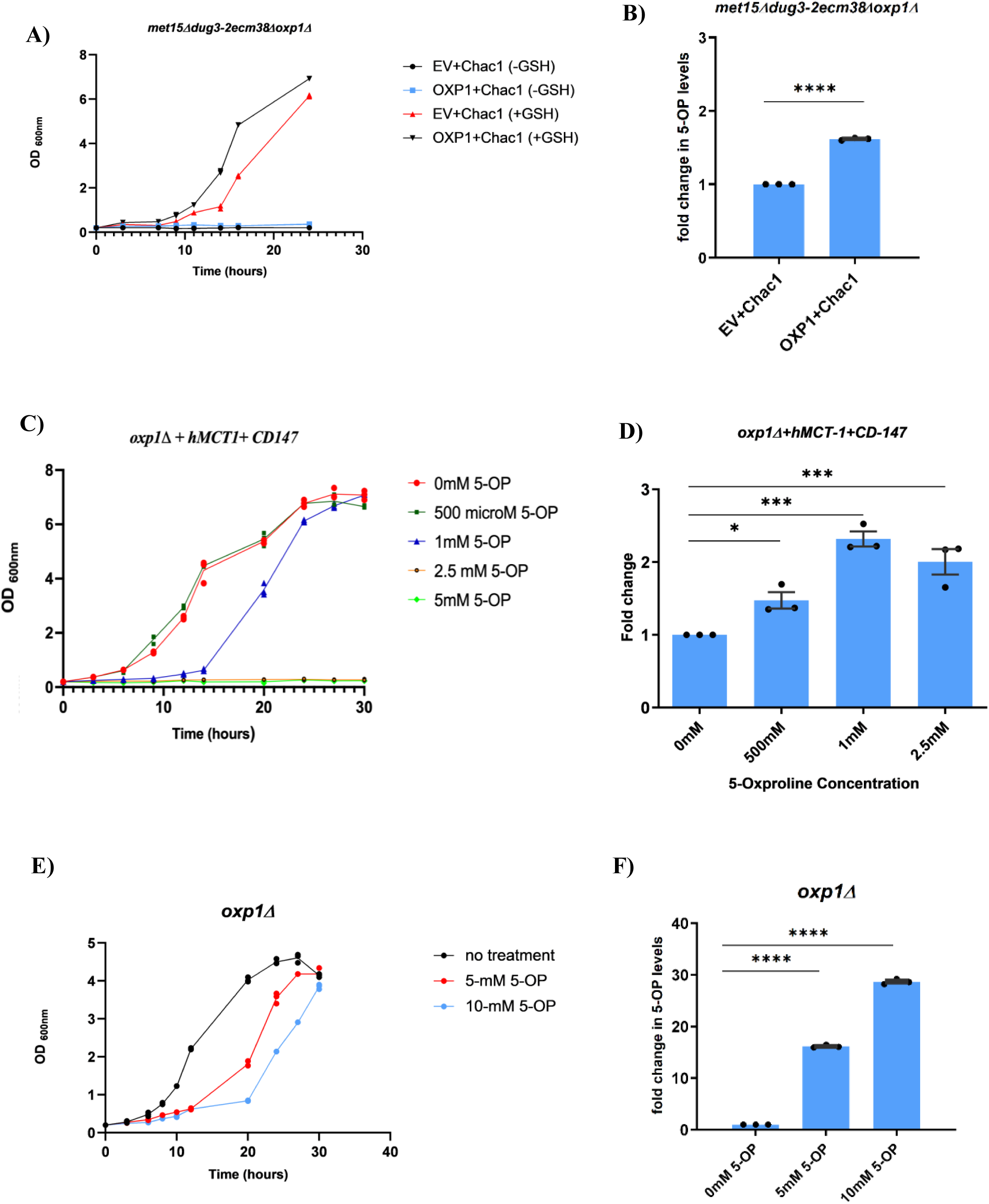
Growth and 5-oxoproline estimation in different models and its consequences: A) Growth curve analysis comparing human Chac1-overexpressing cells carrying either empty vector or co-expressing TEF-OXP1. Only a mild growth difference was observed. B) Quantification of 5-oxoproline (5-OP) in Chac1-overexpressing cells co-expressing either OXP1 or empty vector. Samples were collected and 5-OP estimated as described in Materials and Methods. Statistical analysis was carried out using Student’s t-test. Significance is indicated as (*p < 0.05, **p < 0.01, ***p < 0.001, ****p < 0.0001). error bars represent SD and SEM on growth plots and bar plots respectively. C) Growth of cells treated with 5 mM or 10 mM 5-OP. D) Intracellular 5-OP levels in cells treated with 5 mM or 10 mM 5-OP compared to untreated control, quantified by mass spectrometry. Statistical analysis was carried out using one way ANOVA. Significance is indicated as (*p < 0.05, **p < 0.01, ***p < 0.001, ****p < 0.0001). error bars represent SD and SEM on growth plots and bar plots respectively. E) Growth analysis of *oxp1Δ* cells transformed with MCT-1 and CD-147 on lower concentrations of 5-OP (500 μM, 1 mM, and 2.5 mM). Cells grown on 1 mM 5-OP showed a significantly defective growth phenotype and were selected for further studies. F) Mass spectrometry-based quantification of 5-OP in MCT-1/CD-147-transformed *oxp1Δ* cells exposed to 1 mM and 2.5 mM 5-OP, showing significantly elevated 5-OP levels. All experiments were performed independently three times. Statistical analysis was carried out using one way ANOVA. Significance is indicated as (*p < 0.05, **p < 0.01, ***p < 0.001, ****p < 0.0001). error bars represent SD and SEM on growth plots and bar plots respectively.

In the second approach we examined the use of exogenous 5-OP to increase 5-OP levels in the cell. Although in mammalian cells a 5-OP transporter has been described (SLC16A1/MCT1) (Sasaki et al., 2015), in yeast the transporter of 5-OP is not known. We therefore initially evaluated whether heterologous expression of the human 5-OP transporter, MCT1 would enhance the sensitivity of yeasts to 5-OP. MCT1 in human is a pyruvate transporter that was also shown to transport 5-OP (Sasaki et al., 2015). Its expression in yeast was shown to permit transport of pyruvate in yeast but the activity to transport 5-OP was not examined (Makuc et al., 2004). The surface expression of MCT1 was enhanced with expression of CD-147, although even then only 3% was surface localized (Kirk et al., 2000). The ability of the heterologously expressed MCT1 under the TEF promoter to enhance 5-OP sensitivity in yeast was thus examined (with and without CD147). We observed that in these cells, the sensitivity to 5-OP was significantly enhanced since even at 1mM 5-OP the growth of cells was retarded in these transformants (Fig1E). When the cells only contained hMCT1, there was some growth retardation in the presence of 5-OP but this was enhanced when MCT1 and CD147 were both expressed together (Fig S1). We quantified 5-OP levels and there was a significant accumulation observed even at 1mM concentration (Fig1F). These results indicate firstly that human MCT1 can transport 5-OP in yeast, and secondly that the growth retardation of yeast seen in the presence of 5-OP is a consequence of its transport into the cell and not due to its action on the surface membrane. However, rather than using the heterologously expressed proteins which had reportedly limited surface localization, we decided to resort to the yeast model without any heterologously expressed proteins to simplify the subsequent downstream conclusions.

We evaluated increasing amount of 5-OP in the media to *oxp1Δ* cells and looked for concentrations at which growth defects were seen (Fig 1C). In liquid medium we found that beginning with 5mM, and more significantly at 10mM we could see a retardation in growth and we subsequently used these concentrations. We then estimated 5-OP levels in cells treated with either 5mM 5-OP or 10mM 5-OP after extensively washing off any surface 5-OP. We observed that in the 5mM 5-OP-treated cells the intracellular levels of 5-OP increased 12-fold, while in 10mM it had increased 20-fold (Fig 1D). These elevated levels seemed sufficient to evaluate the consequences on the cell.

### Transcriptomic analysis to study the consequences of 5-OP accumulation

To obtain insights into the consequences of cellular treatment of 5-OP and its accumulation in yeast we carried out a transcriptome sequencing of yeast cells exposed to 5mM 5-oxoproline. We chose 5mM because at this concentration, intracellular 5-OP levels were significantly elevated (∼12 fold) and it was beginning to retard the growth of the cells. The transcriptomic approach was done under these two conditions so that we would obtain an unbiased insight into (a) the possible pathways or proteins that were targets of 5-OP leading to their inactivation and consequently the effect on growth and (b) the cellular response to 5-OP toxicity enabling the cells to tackle the 5-OP treatment and accumulation. For these experiments we used *S. cerevisiae oxp1Δ* cells with or without 5-OP treatment. Cells were treated with 5mM (or no treatment) and harvested at log phase of growth followed by RNA isolation and RNA seq analysis using the Illumina platform. We analysed the data after removing those with insignificant *p* Value (>0.05), and carried out a GO enrichment analysis using the ShinyGO algorithm (Reference). We observed, while using cutoffs of E<0.05 and Fold enrichment >10, that among the biological processes only the xenobiotic export, xenobiotic detoxification and export across the plasma membrane (Supplementary Table S3). All these pathways largely included the same subset of genes, although it missed one in the class, FEX2 that we picked up manually. Among the downregulated genes, using the same cutoffs, no biological processes appeared as significant in the ShinyGO analysis (Supplementary Table S3). Among the upregulated genes, genes of the oxidative stress pathway seemed particularly absent, instead we found many genes that were 2-fold induced. and manual observations also confirmed that these were in multiple categories or pathways (Table 1), We used a cut off of log_2_fold =1.20. The upregulated include the efflux pumps (PDR5, ERC1, SNQ2, FEX2), membrane transporters (PMA1, PNS1, YLR413w, PRM10, WSC2, FLC2), transcription factors (HCM1, SKN7, SMP1, TBF1), enzymes (KDX1, DBP8, HMT1, AAA1, IME2, YHR033W). However there did not seem any particular pathway that was primarily affected other than the xenobiotic export or efflux pathways.

**Table 1:** List of upregulated genes; transcriptomics analysis was done between WT and *oxp1Δ* cells grown on 5mM 5-OP at 2.5 OD.

| Gene | log2FoldChange | pvalue | Protein Names | Gene ontology |
| --- | --- | --- | --- | --- |
| **YHR033W** | 1.88 | 1.82E-06 | Uncharacterized protein YHR033W | Unknown function |
| **YLL066C** | 1.79 | 0.009487 | Y' element ATP-dependent helicase YLL066C | DNA helicase activity |
| **KDX1** | 1.68 | 3.72E-05 | Serine/threonine-protein kinase KDX1 | Protein phosphorylation |
| **PRM10** | 1.66 | 0.002075 | Pheromone-regulated membrane protein 10 | Response to pheromone |
| **INA1** | 1.54 | 0.000328 | Cell membrane protein YLR413W | Cell adhesion |
| **ELG1** | 1.44 | 0.00318 | Telomere length regulation protein ELG1 | DNA replication |
| **RRT5** | 1.44 | 0.170165 | Regulator of rDNA transcription protein 5 | rDNA transcription regulation |
| **FEX2** | 1.44 | 0.017223 | Fluoride export protein 2 | Fluoride ion transport |
| **ERC1** | 1.43 | 0.011175 | Ethionine resistance-conferring protein 1 | Response to toxic substance (ethionine) |
| **PDR5** | 1.41 | 0.000132 | Pleiotropic ABC efflux transporter of multiple drugs | Drug export; ATPase activity |
| **PMA1** | 1.41 | 0.058436 | Plasma membrane ATPase 1 | Proton transport; ATP hydrolysis |
| **OSI1** | 1.38 | 0.029716 | Uncharacterized oxidoreductase YKL071W | Oxidoreductase activity; stress response |
| **PET122** | 1.37 | 0.006165 | Protein PET122, mitochondrial | Translation activator |
| **TBF1** | 1.35 | 8.35E-05 | Protein TBF1 | DNA binding; transcription regulation |
| **PLM2** | 1.34 | 0.005771 | Protein PLM2 | Unknown function |
| **SMP1** | 1.34 | 0.010139 | Transcription factor SMP1 | Transcription regulation; MAPK signaling; stress response |
| **PNS1** | 1.31 | 0.025734 | Protein PNS1 (pH nine-sensitive protein 1) | Vacuolar acidification |
| **HCM1** | 1.30 | 0.000112 | Forkhead transcription factor HCM1 | Transcription regulation |
| **DBP8** | 1.30 | 0.003044 | ATP-dependent RNA helicase DBP8 | rRNA processing |
| **SWE1** | 1.28 | 0.000211 | Mitosis inhibitor protein kinase SWE1 | Cell cycle checkpoint |
| **SKN7** | 1.27 | 0.014344 | Transcription factor SKN7 | Oxidative stress response |
| **HMT1** | 1.25 | 0.002766 | Protein arginine N-methyltransferase 1 | Protein methylation |
| **AAH1** | 1.24 | 0.000695 | Adenine deaminase (ADE) | Purine metabolism |
| **PHM7** | 1.23 | 0.011321 | Phosphate metabolism protein 7 | Phosphate ion homeostasis |
| **WSC2** | 1.23 | 0.004092 | Cell wall integrity and stress response component 2 | Cell wall organization; stress response signaling |
| **FLC2** | 1.23 | 0.000806 | Flavin carrier protein 2 (FAD transporter 2) | FAD transmembrane transport; |
| **RET1** | 1.23 | 0.000607 | DNA-directed RNA polymerase III subunit RPC2 | Transcription by RNA polymerase III |
| **SNQ2** | 1.21 | 0.002116 | Protein SNQ2 | Multidrug resistance |
| **IME2** | 1.21 | 0.067645 | Meiosis induction protein kinase IME2/SME1 | Meiotic cell cycle; protein phosphorylation |

### Validation of the RNA seq data by evaluation of a few genes by q-RT-PCR

To validate the accuracy of our RNA-Seq data, we selected representative genes from different pathways for further analysis. Specifically, we examined three efflux pumps (PDR5, SNQ2, FEX2) and the transcription factor SKN7. All four genes showed upregulation in response to 5-OP accumulation (Fig S2).

### Evaluation of selected upregulated genes by either overexpression or deletion analysis

In the absence of any prior information about the targets of 5-OP and the response to excess 5-OP, we procured the deletions of a randomly selected list of upregulated genes from Euroscarf and examined them on plates/liquid medium containing higher levels of 5-OP to examine if any of the deletions showed enhanced sensitivity. This was carried out by comparing growth of these strains in liquid medium in the presence of 5-oxoproline (5mM) as well as on plate (100mM). We deal with the different genes category wise.

### Efflux pumps of multiple pathways seem involved in the cellular response to 5-OP

The RNA seq data revealed 4 efflux transporters that were upregulated. Interestingly these 4 efflux proteins fell into three different type of transporters- the ABC transporter (PDR5, SNQ2), the multidrug/oligo-saccharide-lipid (polysaccharide (MOP) exporter family (ERC1) and the fluoride exporter (FEX) family. PDR5 and SNQ2, are known to represent functional and structural homologues of mammalian MDR1 and MRP transporter (Zaman et al., 1994). Transcription of both PDR5 and SNQ2 in yeast is controlled by the transcription regulatory proteins Pdr1p and Pdr3p (Balzi et al., 1994; Decottignies et al., 1995). ERC1 gene was primarily identified as a gene conferring ethionine resistance in *S. cerevisiae* and over expression of the ERC1 in *S. cerevisiae* was earlier shown to lead to SAM accumulation (Shiomi et al., 1991). FEX2 is involved in fluoride export and part of a widespread family of conserved fluoride export proteins. FEX2 is a paralog of FEX1, and deletion of both proteins results in a large increase in fluoride sensitivity compared with the single mutant in yeast. It contains two FEX domains connected by a linker (Li et al., 2013). The upregulation of three of these efflux transporter genes (PDR5, SNQ2, FEX2) was validated through q-RT-PCR as described earlier. The deletion of these strains was sensitive on plate containing 100mM 5-OP, suggesting their involvement in efflux of 5-OP (Fig 2A) The growth was also confirmed in liquid media (Fig S6). To further confirm their involvement in 5-OP efflux, the genes were cloned downstream of the strong TEF promoter. The overexpression of PDR5, SNQ2, ERC1 genes was seen to provide resistance to high concentrations of 5-OP on plate which further confirms the involvement of these efflux pumps in rescuing the cell from toxic effect of high amounts of 5-OP in *oxp1Δ* cells (Fig 2B). Only overexpression of FEX2 did not confer resistance although we did not check if this was due to inadequate Fex2p expression. However, the deletion of *fex2Δ* was sensitive. Overexpression of PDR5 rescued the defective phenotype of *pdr5Δ* (Fig S2). Overall, this data suggests that these four efflux pumps of different families are all involved in the rescue of the cells from 5-OP toxicity. These efflux pumps are known to export wide range of compounds and our data suggests that 5-OP is also one of the substrates exported out by these pumps.

**Figure 2.**
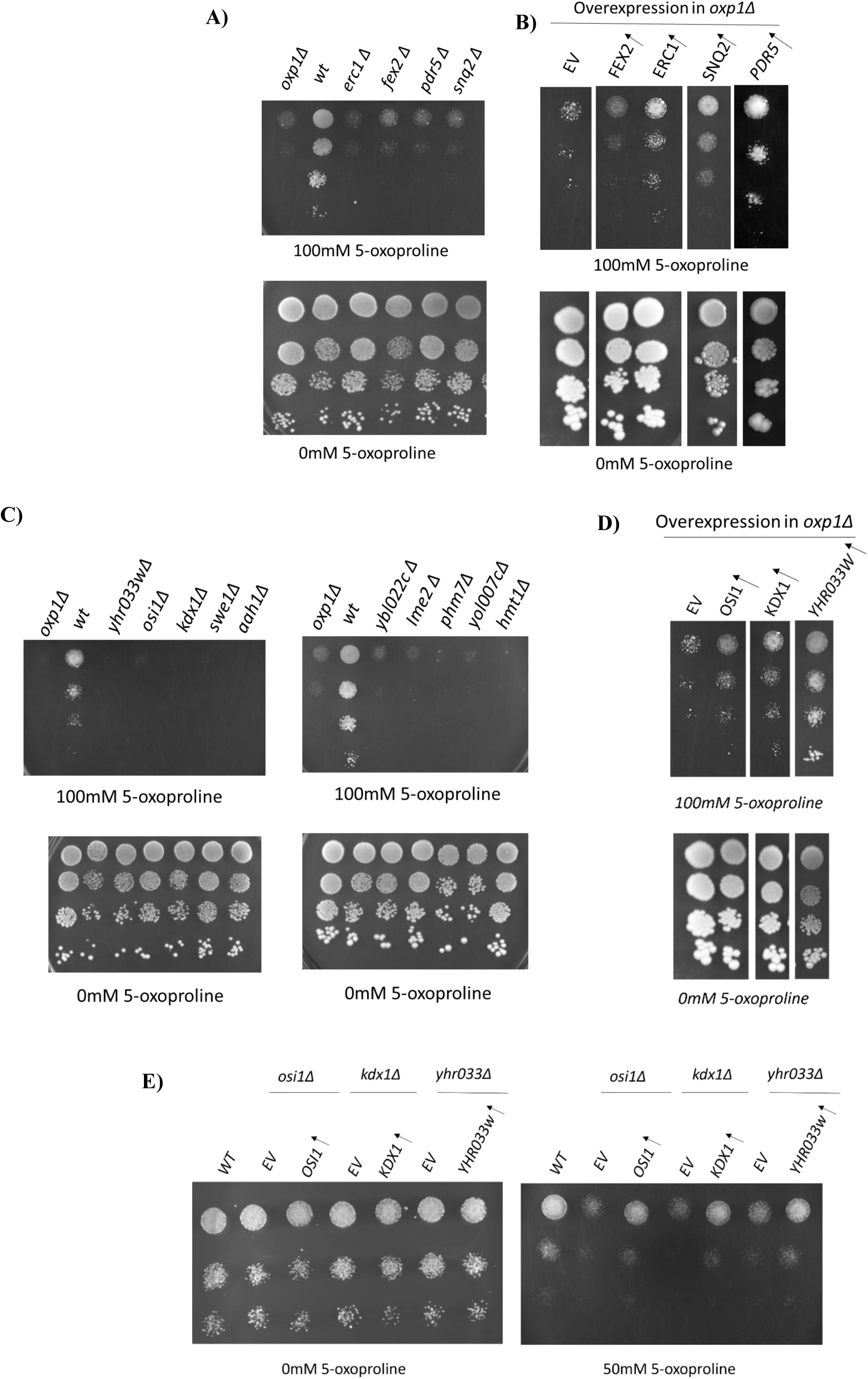
Involvement of efflux pumps and metabolic enzymes in 5-oxoproline (5-OP) resistance: A) efflux deletion strains were sensitive to 5-OP similar to *oxp1Δ*, compared to WT (B) Overexpression of upregulated efflux pumps in *oxp1Δ* cells to evaluate resistance conferred on them. WT cells were BY4741 (AB5000), *oxp1Δ* was in BY4741 background C) Evaluation of different enzymes deletion strains on 5-oxoproline, strains were compared with *oxp1Δ* and the WT (BY4741). **D) O**verexpression of OSI1, KDX1 and YHR033W genes in *oxp1Δ* cells confers resistance towards 5-OP. Images were capture on day 4. E) **O**verexpression of OSI1, KDX1 and YHR033W genes in their respective Δ cells confers resistance towards 5-OP. All experiments were performed independently three times, data shown are representative images.

### Enzymes in different pathways appears to be targets of 5-OP toxicity

A variety of different enzymes in different pathways were found to be upregulated during 5-OP exposure. This also included a few potential enzymes of uncharacterized function. YHR033W encodes a putative glutamate 5-kinase and a paralog of PRO1 protein but no experimental evidences are present for its kinase activity. OSI1 encode a short-chain dehydrogenase/reductase (SDR) protein with NADH dependent enzymatic activities for reduction of furfural (FF), glycolaldehyde (GA), formaldehyde (FA), and benzaldehyde (BZA) (Wang et al., 2017). This protein found to be upregulated under glycolaldehyde (GA) and furfural stress conditions (Wang *et al.,* 2017). KDX1 encodes a Protein kinase that is implicated in Slt2p mitogen-activated (MAP) kinase signalling pathway. It is also known as a stress-responsive protein and interacts with Rlm1p to activate RCK1 gene expression in response to stress in *S. cerevisiae* (Chang et al., 2013). As some of these could be potential target of the accumulating 5-OP we evaluated deletions of these strains for any alterations in 5-OP toxicity (Fig 2C) and a few in liquid cultures (Fig S6). The deletion strains for genes encoding these enzymes were sensitive to 5-OP while overexpression of YHR033w, KDX1 and OSI1 confers a resistance to 5-OP toxicity in *oxp1Δ* strain as well as their respective deletion strains (Fig 2D, E). This data suggests that high levels of 5-OP interfere with proper functioning of different metabolic pathways in the cell and disturbs the cellular homeostasis and results in defective growth.

### Investigating the role of different DNA binding proteins and transcription factors

The transcriptomics data revealed a few DNA binding proteins and transcription factors acting in diverse pathways to be upregulated. These include *SKN7, ELG1, RRT5, TBF1, SMP1* and *HCM1*. We evaluated the deletion of the various transcription factors. We observed that many of the transcription factor deletions showed a mild sensitivity to 5-OP, but the maximum sensitivity was seen with *skn7Δ*. Since 5-OP has been reported to cause oxidative stress and one of the stress response factors Skn7p was upregulated we examined this in greater detail. Its involvement was confirmed through the plate assay where the *skn7Δ* strain was sensitive to 5-OP on plate containing 100mM 5-OP (Fig 3A). It was also evaluated through liquid culture. Skn7p, is a transcription factor which works under stress condition and also has a role in oxidative stress response in association with the Yap1p transcription factor (Lee et al., 1999). Overexpression of Skn7 in *oxp1Δ* strain provides resistance towards 5-OP toxicity (Fig 3B). However, overexpression of YAP1 shows barely any resistance to 5-OP compared to SKN7overexpression (Fig 3B). Other oxidative stress response transcription factor deletion strains such as *rrt5Δ, hcm1Δ, smp1Δ*, *msn2Δ, msn4Δ* were not showing sensitivity towards 5-OP except *elg1Δ* and *tbf1Δ* showing a mild sensitivity (Fig S4).

**Figure 3.**
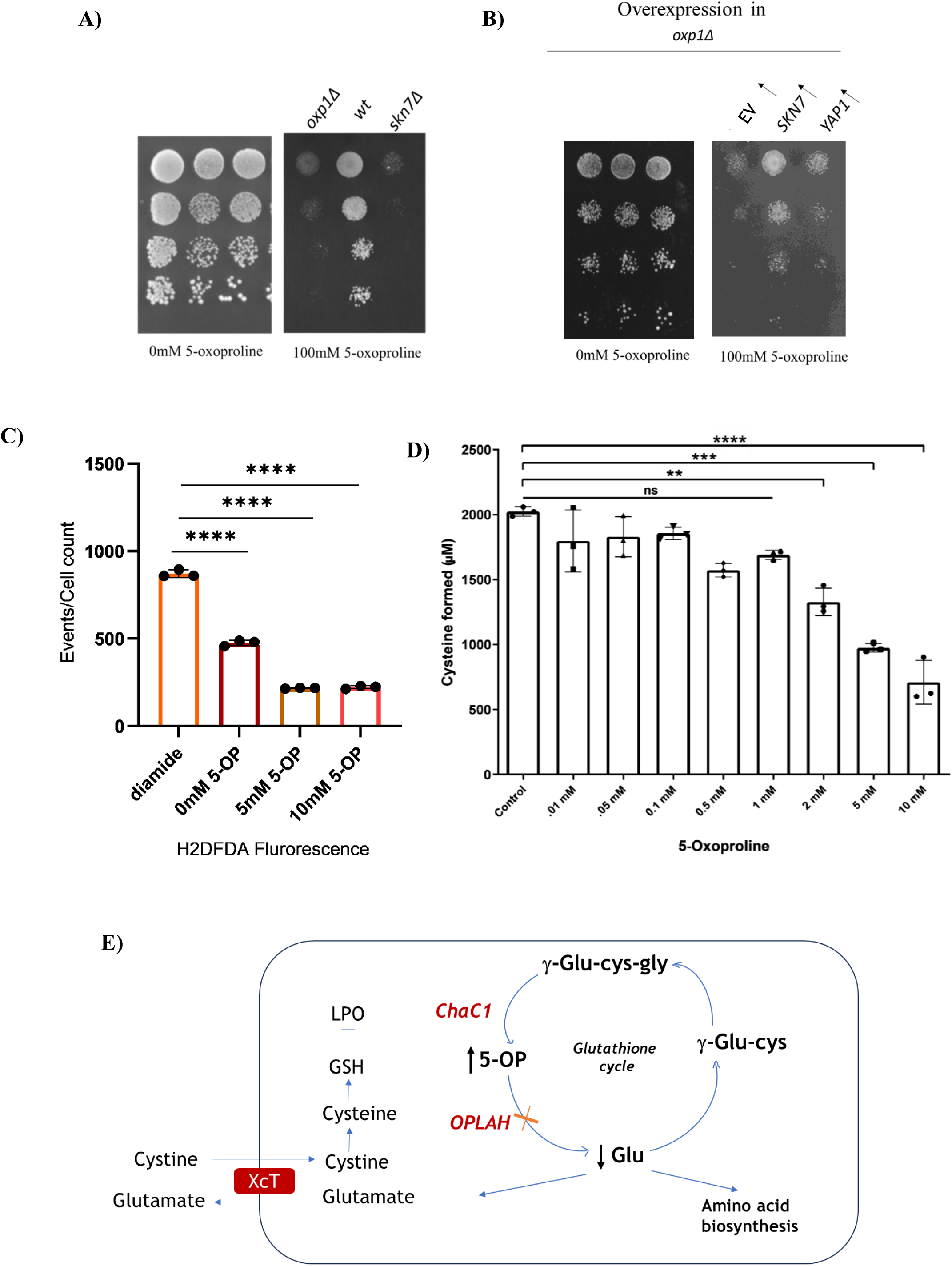
Stress response and Chac1 inhibition by 5-oxoproline (5-OP): A) *skn7Δ* strain showed sensitivity to 100 mM 5-OP compared with WT (BY4741). B) Overexpression of SKN7 conferred strong resistance to 5-OP, whereas YAP1 overexpression resulted in mild resistance. Images were capture by day 4. C) ROS measurement by FACS using H2CDFDA dye. Cells grown with 1 mM or 2.5 mM 5-OP for 30 min were incubated with H2CDFDA; diamide treatment was used as a positive control. D) In vitro inhibition of recombinant human Chac1 by increasing concentrations of 5-OP, with activity measured as described in Materials and Methods. All experiments were performed independently three times. Representative images are shown for the spotting assay. Statistical analysis was carried out using One way ANOVA. Significance is indicated as (*p < 0.05, **p < 0.01, ***p < 0.001, ****p < 0.0001), error bars represent SD. E) Schematics showing an alternate hypothesis for 5-OP mediated oxidative stress in mammalian system.

### 5-OP accumulation affects various membrane proteins in the cell

Among the list of upregulated genes, we have found the upregulation of a few membrane proteins upon 5-OP exposure. This includes proteins with functions in cell wall integrity as well as some with unknown functions. In budding yeast, the cell wall integrity pathway is activated by several membrane proteins like WSC2 that act as sensors. These sensor proteins activate Pkc1, which in turn activates the MAPK module by phosphorylation (Chen and Thorner, 2007). The *S. cerevisiae* FLC2 protein is a putative calcium channel with a role in cell wall integrity, FLC2 gene deletion resulted in pleiotropic phenotypes (Protchenko et al., 2006). PRM10 is selectively expressed during mating and that encodes a multi-spanning transmembrane protein. It has a role in plasma membrane fusion and localizes to sites of cell–cell contact where fusion occurs (Heiman and Walter, 2000). In our study we observed that deletion of these genes shows sensitivity to 5-OP on plate (Fig S5) but the exact mechanisms would need further exploration.

### Evaluation of oxidative stress response in 5-OP treated yeast cells

Since Skn7p, a stress response transcription factor showed sensitivity to 5-OP and since 5-OP was suggested to cause oxidative stress in cardiac tissue, we decided to evaluate the oxidative stress. We evaluated the presence of ROS in the cells using H2CDFDA dye. We did not observe any increase in ROS levels in the 5-OP treated cells compared to diamide treated cells which were used as a positive control (Fig 3C). This result suggests that 5-oxoproline accumulation do not cause ROS dependent oxidative stress.

### 5-OP accumulation does not inhibit the Chac1 enzyme

The accumulation of 5-OP induces a general stress response in cells, yet the initial steps triggering this response remain unclear, as 5-OP itself appears an inert molecule. Further in human cells, the 5-OP accumulation was also accompanied by an oxidative stress response. The human Chac1, an efficient degradation of glutathione with one of the products being 5-OP. There, there was a possibility that 5-OP accumulation might be exhibiting feedback regulation of Chac1. We therefore investigated the effect of 5-OP on the human Chac1 under *in vitro* conditions. We observed a very mild inhibition.This also was observed only at higher concentrations of 5-OP (10mM) (Fig 3D). Thus, it does not appear that Chac1 is a target of the elevated 5-OP levels.

## Discussion

5-oxoproline, as a metabolite, has been known for a long time. However, it has only been actively investigated as an N-terminal moiety of proteins. As a metabolite in cells there have been limited studies. Following the observations that 5-oxoprolinase (OPLAH) knockdown have higher propensity for heart failure in mouse and man has brought the substrate of OPLAH, 5-oxoproline, into focus. To understand how higher 5-OP in the cells affect cell metabolism we needed a suitable model. The well-defined yeast *S. cerevisiae* appeared to be an excellent eukaryotic model to investigate the consequences of 5-OP. The ideal model of 5-OP accumulation would have been one where the 5-OP was generated endogenously. However, this approach using human Chac1 did not succeed in generating elevated levels of 5-OP. It might be because the levels of GSH in the cell were not high enough. We eventually succeeded in creating a model that had between 12- to 20- fold higher 5-OP levels, substantially higher folds than observed in mammalian system.

Investigations using this model revealed that elevated 5-oxoproline is relatively well tolerated by cells. There was only a mild upregulation of genes of diverse pathway. The second key finding from this study is that 5-OP does not cause oxidative stress. This is seen from both the transcriptomics studies and the ROS determination. However, the study did not test the changes in protein levels. Since 5-OP is primarily formed from the degradation of GSH by Chac1, and since this has an indirect effect on the redox environment, we also checked if Chac1 might be feedback regulated by one of its products (5-OP). Glutathione concentrations in the cytoplasm are typically in the range of 1-2mM (although in some cells such as hepatocytes it can be upto 10mM) (Forman et al., 2009). Concentrations of glutathione around 2mM is also consistent with the Km of ChaC1 for glutathione (which is 2.2 mM) (Kaur et al., 2017). Given the concentrations of glutathione in the range of 2mM, the concentrations of 5-oxoproline (the product of the reaction with ChaC1) are unlikely to exceed 2mM. However, with 2mM 5-oxoproline only 33% inhibition was observed. Even at 10mM of 5-oxproline only 65% inhibition was observed. This limited inhibition of Chac1 cannot explain the severe downstream effects seen in cardiac cells even though it could be a partial contributor. Indeed, when we examine the cardiac study carefully, we see that the authors had found only 2-fold increase in plasma level of 5-oxoproline in OPLAH knockdown. This is not a significant increase because 5-OP levels in plasma vary several folds only with dietary differences (Jackson et al., 1996; Metges et al., 2000). One possible criticism of this study or that while what we are observing with yeast may not be true, in case of mammalian cells response could be different and one might be experiencing oxidative stress response in the mammalian system, though not the yeast model. However, the observations are consistent with the earlier suggested role of 5-oxoproline as a relatively inert with a role as an osmoprotection.

If 5-oxoproline (5-OP) levels are not the direct cause of the observed oxidative stress and phenotypic consequences, then what might be the possible explanation? One plausible alternative is that the downstream metabolite of 5-OP, glutamate, produced by the enzyme 5-oxoprolinase (OPLAH), plays a critical role. Glutamate could contribute to maintaining redox balance and cellular homeostasis by enhancing cysteine availability through the cystine/glutamate antiporter (XcT). This exchange mechanism imports cystine in exchange for glutamate, and once inside the cell, cystine is reduced to cysteine, a rate-limiting precursor for glutathione (GSH) synthesis. Therefore, disruptions in 5-OP metabolism could indirectly affect glutathione biosynthesis and redox buffering capacity by altering intracellular glutamate and cysteine levels. Additionally, glutamate serves as a key amino acid in multiple metabolic pathways, and its altered availability might influence cellular energy metabolism, amino acid balance, or signalling processes that collectively contribute to the observed phenotypes. Importantly, since both glutamate export and cysteine import via system XcT are tightly linked to ferroptosis regulation, changes in glutamate or cysteine availability resulting from altered 5-OP metabolism could significantly impact cellular susceptibility to oxidative stress (Figure 3E). More studies would be needed to confirm the validity of this possibility. However, as yeast lack XcT, it would have to be evaluated in a mammalian model.

## Materials and methods

### Chemicals and Reagents

All chemicals used were of analytical reagent grade. Media components were purchased from Hi Media (India), Merck (Germany) and BD Difco (USA). Oligonucleotides were purchased from Merck (Germany) and GCC (India). Restriction enzymes, Phusion polymerase, dNTPs and other modifying enzymes were obtained from New England Biolabs (Beverly, MA, USA). Gel-extraction kits and plasmid miniprep columns were obtained from QIAGEN (Germany) or Thermo-Fischer Scientific (USA). 5-oxoproline and glutathione were purchased from Merck (Germany).

### Strains, media and Growth

The *Escherichia coli* strain DH5α was used as a cloning host. The list of yeast strains used in this study is shown in Supplementary Table 1. Yeasts were routinely maintained on YPD medium. The minimal medium contained YNB, glucose, and ammonium sulfate supplemented with the required amino acids and bases. Yeast DNA Isolation and Yeast Transformation—Yeast chromosomal DNA was isolated by the glass bead lysis method and yeast transformations were carried out using the lithium acetate method (Gietz and Woods, 2002).

### Growth studies

The growth assay for the endogenous model for 5-OP accumulation was done with *met15Δdug3-2 ecm38Δoxp1Δ* strains transformed with human Chac1 along with OXP1 (test) and human Chac1 with empty vector (control). The transformants were grown overnight on synthetic medium (without ura and his). Further, the inoculum (25 ml) was initiated at 0.01 OD_600_ on complete synthetic medium (without ura, his and met) supplemented with 100μM glutathione (GSH) and growth was monitored. Cells were harvested at mid logarithmic phase and 5-OP quantification was done using mass spec.

For exogenous model, the *oxp1Δ* cells and *oxp1Δ* cells transformed with MCT-1and CD-147 were grown in complete synthetic medium overnight, WT cells (with or without empty vector) were grown on same condition as a control. Further, the inoculum (25 ml) was initiated at 0.01 OD_600_ on complete synthetic medium supplemented with different concentrations (1mM, 2.5mM, 5mM and 10mM) 5-oxoproline and growth was monitored for 32 hours. A without treatment condition was taken as a control.

### Growth assay by dilution spotting

For growth assay, the different strains were grown overnight in minimal ammonia medium without Uracil/Leucine/histidine and reinoculated in fresh medium to an OD600 of 0.1 without Uracil/Leucine/histidine depending upon the selection marker and grown for 6 hr. The exponential-phase cells were harvested, washed with water, and resuspended in water to an OD600 of 0.2. These were serially diluted to 1:10, 1:100, and 1:1000. Of these cell resuspensions, 10μl were spotted on minimal medium containing 50mM or 100mM 5-oxoproline. The plates were incubated at 30 degrees for 3-6 days and photographs were taken.

### RNA sequencing

To study the consequences of 5-OP accumulation AB6302 strain was grown in SD medium containing 5mM 5-oxoproline (test) or 0mM 5-oxoproline (control). The mid log phase cells were harvested, washed with water and frozen. Total RNA was isolated using Qiagen RNeasy mini kit (Cat # 74106), RNA sequencing libraries were prepared with Illumina-compatible NEBNext® Ultra™ II Directional RNA Library Prep Kit (New England BioLabs, MA, USA) were carried out by Genotypic Pvt.Ltd. Further, paired-end Illumina Next Generation Sequencing was performed at Genotypic Technology Pvt. Ltd., Bangalore, India. The experiment was done in duplicates and the data was deposited in NCBI (Accession no. PRJNA1345586).

### GO Enrichment Analysis

GO enrichment analysis was carried out using web-based ShinyGO version 0.85 (http://bioinformatics.sdstate.edu/go/; Ge et al., 2019). The list of genes either upregulated or downregulated in response to 5-oxoproline were subjected to GO enrichment analysis. top biological processes selected using the e value<0.05 and fold enrichment >10, as described in Supplemental Table S3.

### Gene cloning

KDX1, YHR033W, OSI1, ERC1 and FEX2 genes were cloned in pRS416TEF by PCR amplification followed by homologous recombination in yeast as described previously (Singh et al., 2024). The gene specific primers carried overlapping sequences for pRS416TEF and an internal restriction site BamHI and XhoI were also added in forward and reverse primers respectively (Table S2). Two further sets of primers were also designed to amplify the vector in two different fragments carrying the overlapping sequences for homologous recombination. One fragment of the vector called CEN fragment as it contains CEN another called URA fragment as it contains URA marker. For homologous recombination the CEN and URA fragments along with the SEO1 gene fragment, carrying vector overlapping sequences, were transformed in yeast using lithium acetate method and plated on SD-URA plates. A few colonies were picked up and desired recombinants were confirmed by PCR. The Plasmids were isolated after passage through E. coli and digestion by the restriction enzymes BamHI and XhoI confirmed the presence of the clone. Human MCT-1 and CD-147 cDNA was purchased from Sino biologicals, cloned in pRS416-TEF and pRS315-TEF using restriction digestion method under EcoRI and XhoI sites respectively. Other clones used in this study were previously available in the lab.

### Metabolite extraction

Intracellular metabolites for MS-based targeted metabolomics were extracted using 80% methanol. Briefly, the yeast cell pellet was collected by centrifugation in a microcentrifuge tube and washed three times with sterile water, then quenched with pre-chilled 80% methanol (kept at −80°C), followed by seven cycles of bead beating (using glass beads). The solution was centrifuged at 15000 g at 4□ for 15 minutes, and the supernatant was collected in a fresh tube and dried using a vacuum concentrator operated at room temperature. Extracted metabolites were reconstituted in 150µl of 50% methanol with 0.1% formic acid, centrifuged at 15,000 g for 10 min, and transferred to HPLC vials.

### Mass Spectrometry

The data were acquired using a multiple reaction monitoring (MRM) method on a triple quadrupole hybrid ion trap mass spectrometer (QTRAP 6500+, SCIEX) coupled with an ExionLC UHPLC system (SCIEX). Optimized source and gas parameters were used and data acquisition was performed through Analyst 1.6.3 software in positive ion scan mode. 5 µl of metabolites suspension was loaded and resolved on an Acquity UPLC BEH C18 column (1.7 μm, 2.1 × 100 mm, Waters) using mobile phases of water with 0.1% formic acid (buffer A) and acetonitrile with 0.1% formic acid (buffer B) with a flow rate of 0.3 ml/min and 8 minutes long gradient. The gradient program was employed as follows: buffer B was raised from an initial 2% to 30% in 2.1 minutes. In the next 2 minutes, buffer B was increased to 40% and further to 45% in the next 0.7 minutes. The buffer B concentration was ramped up to 90% in the next 0.1 minutes and kept constant for the next 1.1 minutes. Buffer B concentration was brought to an initial 2% concentration in the next 0.5 and the column was equilibrated at this buffer condition for 1.5 minutes before the next sample injection. Relative quantification was performed using MultiQuantTM software v.3.0 (SCIEX). A pooled QC from samples was used to analyze the technical variability.

### q-RT-PCR

For genes, the primer set listed in Table were used with a final reaction volume of 5μl containing using Maxima SYBR Green qPCR Master Mix (Fermentas, USA). PCR conditions were 95°C for 3 min, then 40 cycles consisting of denaturation at 95°C for 10 s, annealing at 60°C for 10s and extension at 72°C for 30 s, followed by the melting curve protocol with 10s at 95°C and then 60s each at 0.5°C increments between 65°C and 95°C. The reactions were performed in triplicate for each sample. The relative amounts of target gene expression for each sample were calculated using the Livak method formula 2-(ΔΔCT) against an endogenous control actin for genes. Finally, the fold change against the control gene is calculated and plotted. Data analysis was performed by Graph pad prism using a paired t-test. The significance of differences between means were calculated at a 5% level (P< 0.05).

### ROS measurement using FACS

Overnight yeast cultures were inoculated into fresh complete SD medium supplemented with 5-oxoproline and incubated at room temperature until reaching the exponential growth phase. Three replicate cultures were harvested (approximately 3 million cells each) by centrifugation at 6,000g for 10 minutes. The cell pellets were then washed twice with 1X phosphate buffer (0.1 M, pH 7.4). For a positive control, one set of samples was treated with 1.5mM of diamide per 1 mL of sample for one hour at 30-degree C. Subsequently, all samples, including the diamide-treated control and test samples, containing approximately 1 million cells each, were incubated with 250 μL of 80 μM H2DCF-DA prepared in DMSO dye in the same buffer for 20-30 minutes at 30 degree C in the dark. After incubation, the cells were washed with 1X PBS and resuspended in 500 μL of fresh 1X PBS. Finally, cellular fluorescence intensity, an indicator of ROS (reactive oxygen species) production, was measured using either a FACS Aria III Fusion machine using FITC channel.

### In vitro ChaC1p-Dug1p coupled enzymatic assay for inhibition studies

5 ng of recombinantly purified human ChaC1 protein was incubated with the inhibitor compound. All compounds were dissolved in dimethyl sulfoxide (DMSO) and diluted with sterilized distilled water until the concentration of DMSO was 1% for 30 min in a 50μl reaction mixture containing 50 mM Tris-Cl (pH 8.0) and 5 mM DTT. Immediately after, 2 mM of the substrate, glutathione, was added to this mixture and further incubated for 30 min. The enzyme was inactivated by heating at 95□ for 5 min. To this, 10μl of reaction mixture containing 5μg of Dug1p and 20 µM MnCl2 was added and incubated at 37□ for another 1 hour. Cysteine liberated from the above reaction was measured using a ninhydrin-based method (Gaitonde, 1967).

## Statistical analysis

The data has been analysed using student’s t test and One way ANOVA with p value cut off of 0.05 using Graph Pad Prism.

## Data availability

All data are contained within the manuscript. RNA sequencing data was deposited in NCBI (Accession no. PRJNA1345586).

## Supporting information

## Author contribution

PD: Performed most of the experiments, data analysis and manuscript writing; VBS: performed MCT-1 model experiments, data analysis; PS: performed 5-OP quantification and data analysis; SSG: supervised the design and analysis of the 5-OP quantification experiments; AKB: Supervision, experiment designing, conceptualization and manuscript writing.

## Acknowledgement

The work was supported by a grant-in-aid project from the Department of Biotechnology (BT/PR39040/BRB/10/1882/2020) to AKB. PD was a recipient of a Senior Research Fellowship from the Indian Council of Medical Research (ICMR).

