## supplementary for "A yeast model of 5-oxoproline accumulation reveals a general toleration to 5-oxoproline"

Supplementary data:

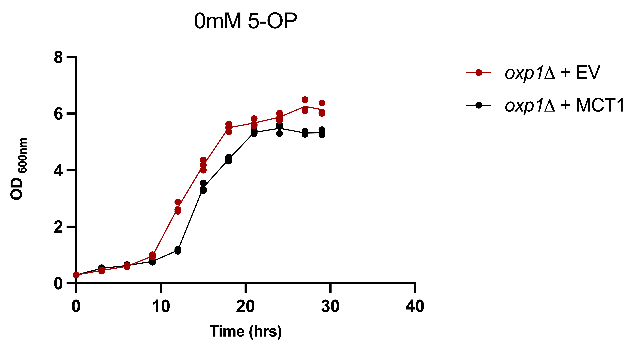

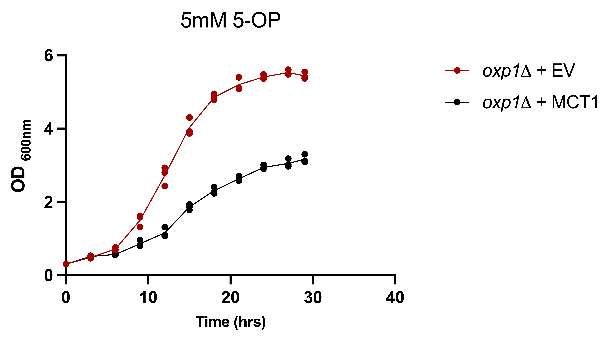

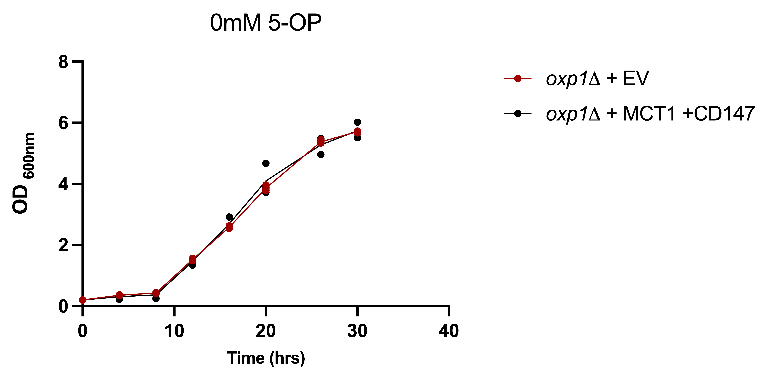

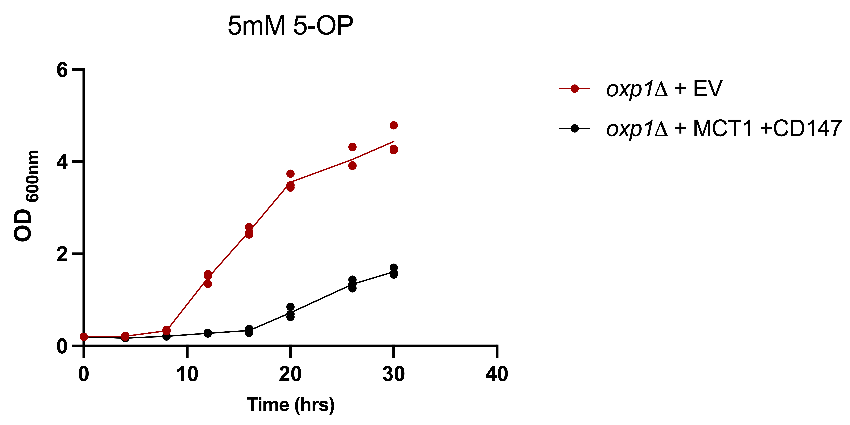

Fig S1: growth pattern of WT and *oxp1Δ* cells with only MCT-1 overexpression as well as along with CD-147, *oxp1Δ* cells with MCT-1 shows severe growth defect on 5mM 5-OP whereas MCT-1 with CD-147 transformed *oxp1Δ* cells were not able to grow.

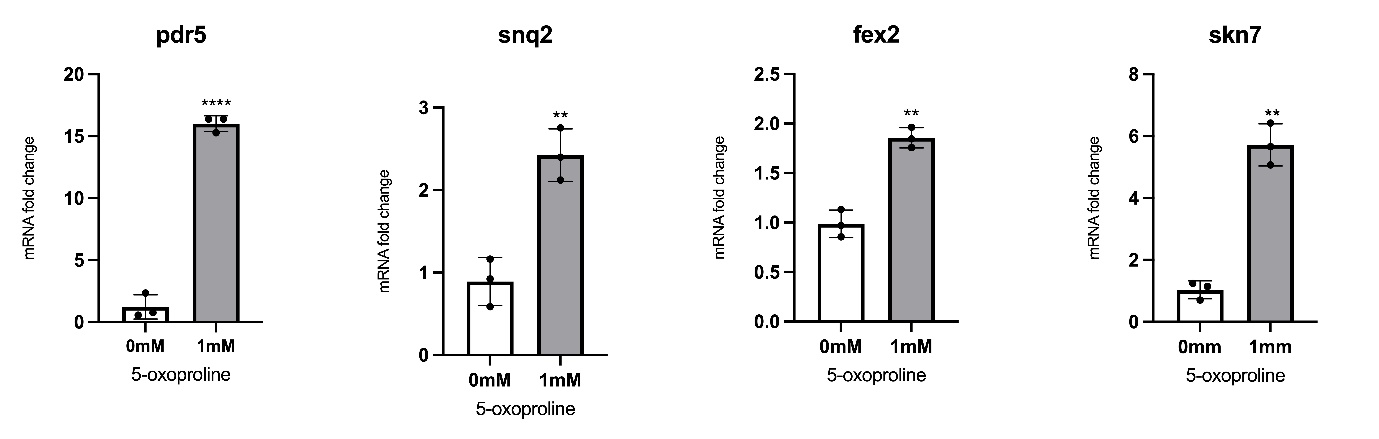

Fig S2: Validation of RNA-seq–identified upregulated genes using qRT-PCR. All experiments were performed independently three times. Statistical analysis was carried out using Student’s t-test. Significance is indicated as (*p < 0.01, **p < 0.001)

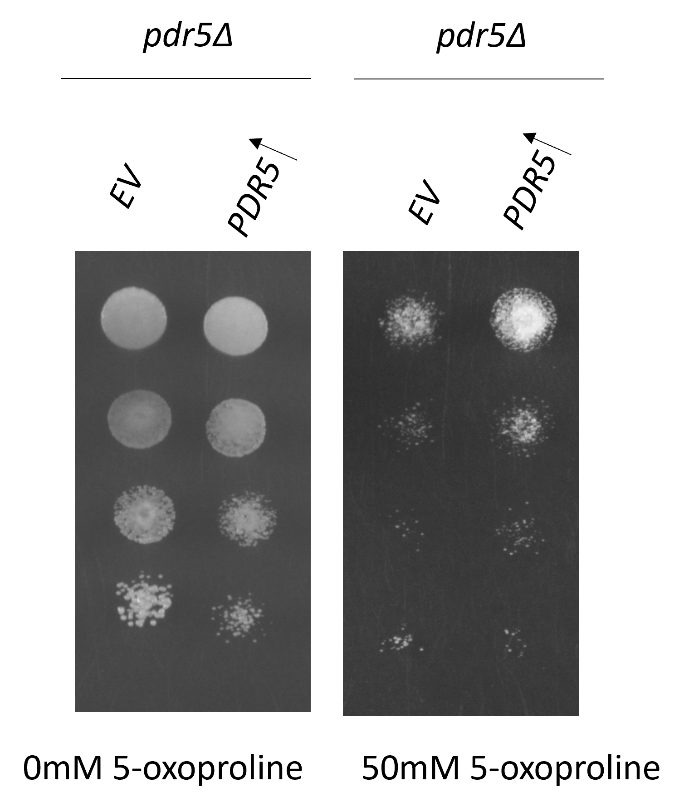

Fig S3: PDR5 overexpression show resistance towards 5-OP in *pdr5Δ* strain, this experiment was repeated to confirm and this is the representative picture. Images were capture by day 4.

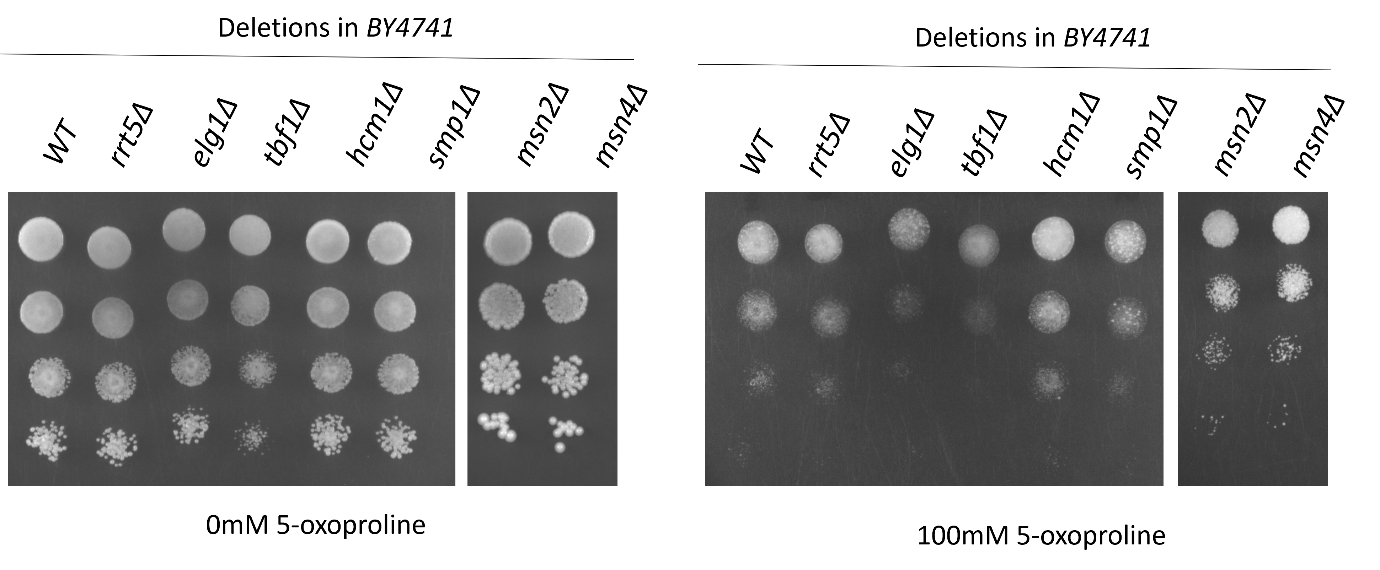

Fig S4: Deletion of other transcription factors did not alter 5-OP sensitivity. Evaluation of membrane protein deletion strains on 5-OP plates compared with WT (BY4741). Images were capture by day 4.

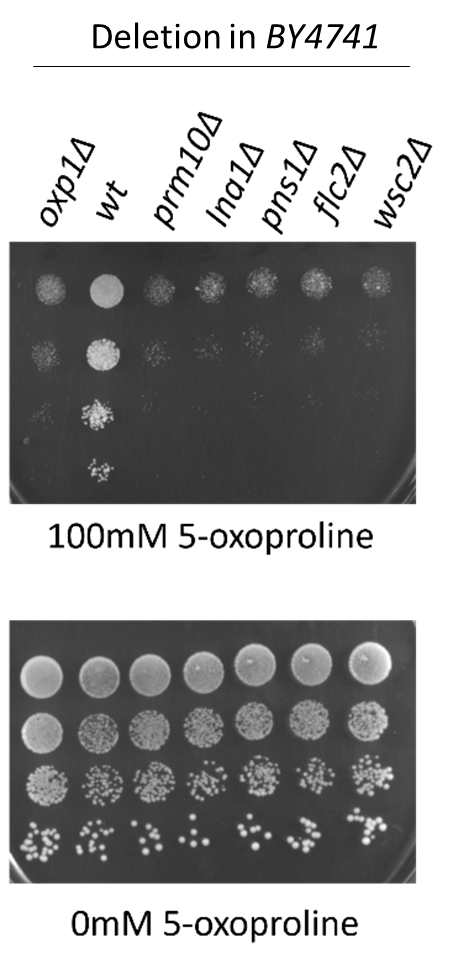

Fig S5: Evaluation of different membrane protein deletion strains on 5-oxoproline, strains were compared with *oxp1Δ* and the WT (BY4741). The deletions strains were sensitive for 5-OP, this experiment was repeated three time and a representative data is presented. Images were capture by day 4.

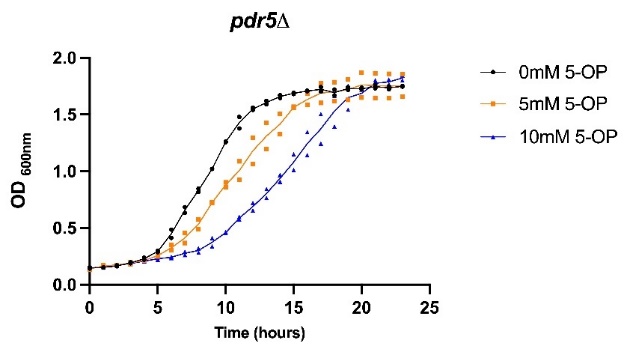

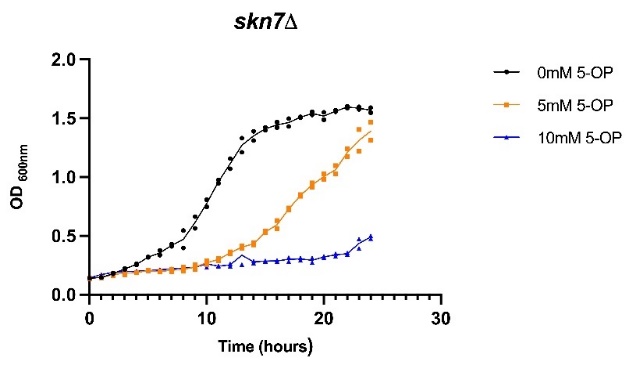

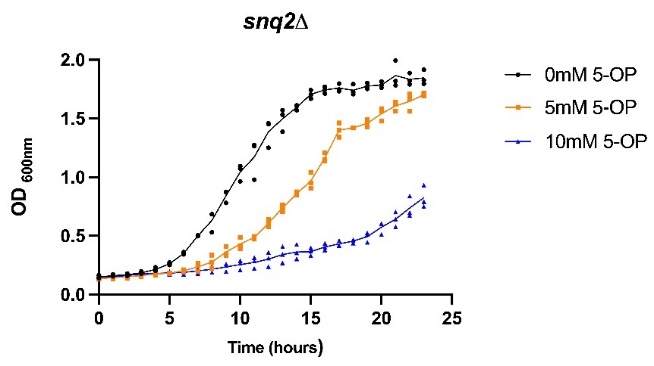

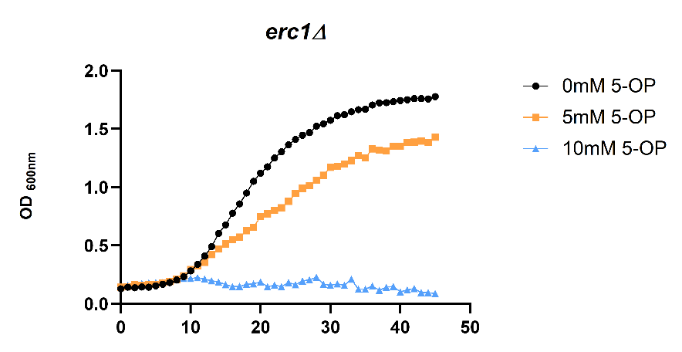

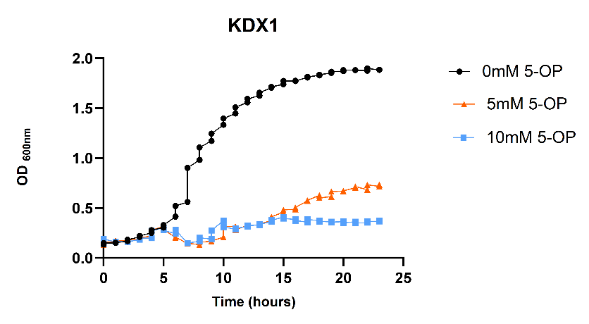

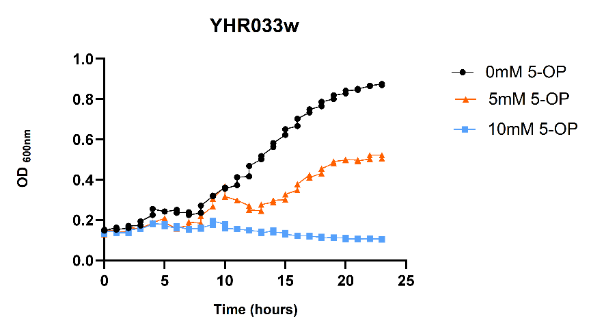

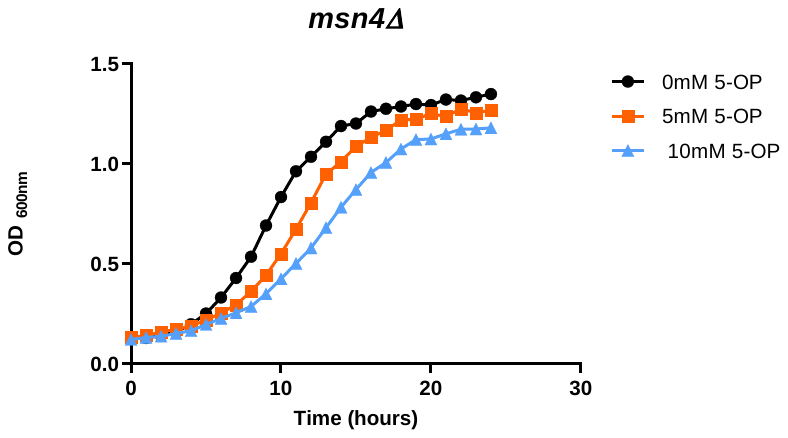

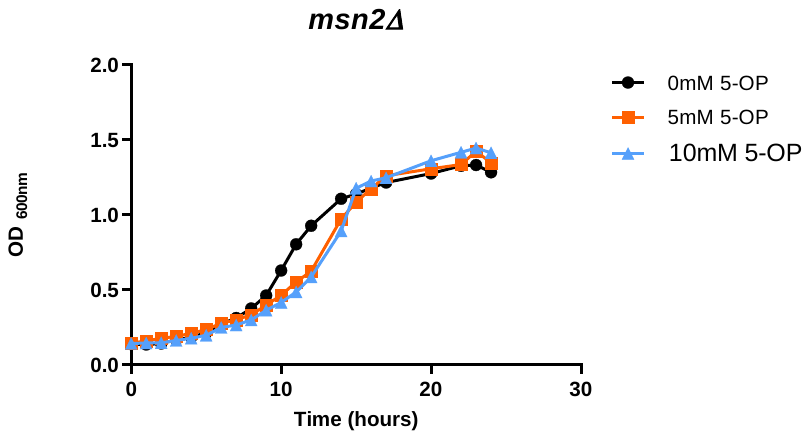

Fig S6: growth analysis of deletion strains in liquid media. All the experiments were done in triplicates, for *pdr5Δ*, *skn7Δ* and *snq2Δ , a*ll three datapoints are shown whereas for *erc1Δ*, *kdx1Δ*, *yhr033wΔ*, *msn4Δ* and *msn2Δ*, a representative is shown.

Supplementary Table1: List of strains used in the study

| Strain | Genotype |
| --- | --- |
| AB5000 | *BY4741MATahis3Δ1leu2 Δ0met15Δ0ura3Δ0* |
| AB6302 | *BY4741MATa his3Δ1leu2Δ 0met15Δ0ura3Δ0oxp1Δ::KanMX* |
| AB1723 | *MATαhis3∆1leu2∆0lys2∆0met15∆0ura3∆0ecm38∆::KanMX4dug3-2* |
| AB6486 | *MATαhis3∆1 leu2∆0 met15∆0 ura3∆0 dug3∆oxp1∆::LEU2* |
| AB6547 | *MATαhis3∆1 leu2∆0 lys2∆0 met15∆0 ura3∆0ecm38∆::KanMX4 dug3-2oxp1∆:: LEU2* |
| AB6259 | *BY4741MATahis3Δ1leu2Δ 0met15Δ 0ura3Δ 0pdr5Δ::KanMX* |
| AB681 | *BY4741MATahis3Δ1leu2Δ 0met15Δ 0ura3Δ 0snq2Δ::KanMX* |
| AB6642 | *BY4741MATahis3Δ1leu2Δ 0met15Δ 0ura3Δ 0erc1Δ::KanMX* |
| AB6634 | *BY4741MATahis3Δ1leu2Δ 0met15Δ 0ura3Δ 0osi1Δ::KanMX* |
| AB6618 | *BY4741MATahis3Δ1leu2Δ0met15Δ0ura3Δ0YHR033wΔ::KanMX* |
| AB6620 | *BY4741MATahis3Δ1leu2Δ 0met15Δ 0ura3Δ 0swe1Δ::KanMX* |
| AB6628 | *BY4741MATahis3Δ1leu2Δ 0met15Δ 0ura3Δ 0aah1Δ::KanMX* |
| AB6629 | *BY4741MATahis3Δ1leu2Δ0met15Δ0ura3Δ 0YBL022cΔ::KanMX* |
| AB6623 | *BY4741MATahis3Δ1leu2Δ 0met15Δ 0ura3Δ 0ime2Δ::KanMX* |
| AB6625 | *BY4741MATahis3Δ1leu2Δ 0met15Δ 0ura3Δ 0phm7Δ::KanMX* |
| AB6624 | *BY4741MATahis3Δ1leu2Δ0met15Δ0ura3Δ 0YOL007cΔ::KanMX* |
| AB6630 | *BY4741MATahis3Δ1leu2Δ0met15Δ0ura3Δ 0hmt1Δ::KanMX* |
| AB1032 | *BY4741MATahis3Δ1leu2Δ0met15Δ0ura3Δ 0skn7Δ::KanMX* |
| AB1034 | *BY4741MATahis3Δ1leu2Δ0met15Δ0ura3Δ 0yap1Δ::KanMX* |
| AB6621 | *BY4741MATahis3Δ1leu2Δ0met15Δ0ura3Δ 0prm10Δ::KanMX* |
| AB6636 | *BY4741MATahis3Δ1leu2Δ0met15Δ0ura3Δ 0ina1Δ::KanMX* |
| AB6627 | *BY4741MATahis3Δ1leu2Δ0met15Δ0ura3Δ 0pns1Δ::KanMX* |
| AB6617 | *BY4741MATahis3Δ1leu2Δ0met15Δ0ura3Δ 0flc2Δ::KanMX* |
| AB6619 | *BY4741MATahis3Δ1leu2Δ0met15Δ0ura3Δ 0wsc2Δ::KanMX* |

Supplementary Table 2: List of primers used in the study

| Primer Name | SEQUENCE (5’-3’) |
| --- | --- |
| RP_416CEN | CTTCTGTTCGGAGATTACCGAATC |
| FP_416CEN | TCTAGAAAACTTAGATTAGATTGC |
| FP_416Ura | TCGCGCGTTTCGGTGATGAC |
| RP_416Ura | CTCGAGTCATGTAATTAGTTATGT |
| Oxp1 del check FP | CAGTAATAATTTTGAGCATTGA |
| Oxp1 del check RP | TGGTTCCGTGTAATTCGATATT |
| FP-hSLC16A1-BamHI | GACCAAGGATCCATGCCACCAGCAGTTGGAG |
| RP-hSLC16A1-XhoI | GATTACCTCGAGTCAGACTGGACTTTCCTC |
| FP CD147-EcoR-I | GATCATGAATTCATGGCGGCTGCGCTGTTC |
| RP CD147-Xho-I | CATTAGCTCGAGTCAGGAAGAGTTCCTCTGG |
| Oxp1 leu FP | ATGCAGAAAGGAAACATAAGAATTGCCATCGATAAGGGTGAACTGTGGGAATACTCAGGT |
| Oxp1 leu RP | ATGCAGAAAGGAAACATAAGAATTGCCATCGATAAGGGTGAACTGTGGGAATACTCAGGT |
| PDR5 FP-RT | CAGTTGCATGAAAGGGGTGC |
| PDR5 RP-RT | CGTTAGCAACACCAACAGCC |
| SKN7 FP-RT | TAAAACCACCGGCGATAGCA |
| SKN7 RP-RT | TCGTGTTCATCAGTCGAGGC |
| SNQ2 FP-RT | CTTCATCCACTTCCGGTGCT |
| SNQ2 RP-RT | AAAGATGCCAGAGTGGAGCC |
| YAP1 FP-RT | TTCCTCCAGCGCTACTTTGG |
| YAP1 RP-RT | AGCCGAGATGGGTTTCTTGG |
| ERC1 FP-RT | TTGAGGGCGAAGAGCAAGAG |
| ERC1 RP-RT | TGTTGCCACGCTTCTTGTTG |
| FEX2 FP-RT | GTGAACTCGTTGAATTCATGCC |
| FEX2 RP-RT | ACTCAAAGTGCCGCAAAATCC |
| KDX1-FP | TTAGAAAGAAAGCATAGCAATCTAATCTAAGTTTTCTAGAGGATCCATGGCGACTGACAC |
| KDX1-RP | GGAGGGCGTGAATGTAAGCGTGACATAACTAATTACATGACTCGAGTTAGTTAACACCCTG |
| OSI-FP | TTAGAAAGAAAGCATAGCAATCTAATCTAAGTTTTCTAGAGGATCCATGAATACTTCATC |
| OSI-RP | GGAGGGCGTGAATGTAAGCGTGACATAACTAATTACATGACTCGAGCTAAAAGACGCCTTC |
| YHR033W-FP | TTAGAAAGAAAGCATAGCAATCTAATCTAAGTTTTCTAGAGGATCCATGACAAAAGCTTA |
| YHR033W-RP | GGAGGGCGTGAATGTAAGCGTGACATAACTAATTACATGACTCGAGTCAAAATTGCGGAG |
| ERC1-FP | TTAGAAAGAAAGCATAGCAATCTAATCTAAGTTTTCTAGAGGATCCATGTCTAAACAATT |
| ERC1-RP | GAGGGCGTGAATGTAAGCGTGACATAACTAATTACATGACTCGAGCTAGTTATACCCAAC |
| FEX2-FP | TTAGAAAGAAAGCATAGCAATCTAATCTAAGTTTTCTAGAGGATCCATGATTTTCAATCC |
| FEX2-RP | GGAGGGCGTGAATGTAAGCGTGACATAACTAATTACATGACTCGAGCTAACAAATCGGGT |

Supplementary Table 3: Gene ontology analysis of RNA seq data, top biological processes selcectd using the *e value<0.05* and fold enrichment >10

Upregulated biological process

| **Enrichment FDR** | **nGenes** | **Pathway Genes** | **Fold Enrichment** | **Pathways (click for details)** |
| --- | --- | --- | --- | --- |
| 4.2E-02 | 3 | 9 | 21.8 | [Xenobiotic export from cell](https://amigo.geneontology.org/amigo/term/GO:0046618) |
| 4.2E-02 | 3 | 9 | 21.8 | [Xenobiotic detoxification by transmembrane export across the plasma membrane](https://amigo.geneontology.org/amigo/term/GO:1990961) |
| 4.4E-03 | 5 | 17 | 19.2 | [Export across plasma membrane](https://amigo.geneontology.org/amigo/term/GO:0140115) |
| 3.4E-02 | 8 | 100 | 5.2 | [Rna methylation](https://amigo.geneontology.org/amigo/term/GO:0001510) |
| 3.4E-02 | 9 | 126 | 4.7 | [Macromolecule methylation](https://amigo.geneontology.org/amigo/term/GO:0043414) |
| 3.4E-02 | 10 | 152 | 4.3 | [Methylation](https://amigo.geneontology.org/amigo/term/GO:0032259) |
| 3.4E-02 | 19 | 498 | 2.5 | [Transmembrane transport](https://amigo.geneontology.org/amigo/term/GO:0055085) |

Downregulated biological process

| **Enrichment FDR** | **nGenes** | **Pathway Genes** | **Fold Enrichment** | **Pathways (click for details)** |
| --- | --- | --- | --- | --- |
| 1.3E-04 | 10 | 18 | 5.8 | [Mitochondrial electron transport cytochrome c to oxygen](https://amigo.geneontology.org/amigo/term/GO:0006123) |
| 5.1E-12 | 27 | 52 | 5.4 | [Mitochondrial respiratory chain complex assembly](https://amigo.geneontology.org/amigo/term/GO:0033108) |
| 1.7E-12 | 29 | 56 | 5.4 | [Mitochondrial respirasome assembly](https://amigo.geneontology.org/amigo/term/GO:0097250) |
| 1.3E-04 | 11 | 22 | 5.2 | [Atp biosynthetic proc.](https://amigo.geneontology.org/amigo/term/GO:0006754) |
| 1.3E-04 | 11 | 22 | 5.2 | [Proton motive force-driven atp synthesis](https://amigo.geneontology.org/amigo/term/GO:0015986) |
| 9.7E-06 | 15 | 32 | 4.9 | [Respiratory chain complex iv assembly](https://amigo.geneontology.org/amigo/term/GO:0008535) |
| 9.7E-06 | 15 | 32 | 4.9 | [Mitochondrial cytochrome c oxidase assembly](https://amigo.geneontology.org/amigo/term/GO:0033617) |
| 1.7E-06 | 19 | 44 | 4.5 | [Cytochrome complex assembly](https://amigo.geneontology.org/amigo/term/GO:0017004) |
| 8.9E-06 | 18 | 44 | 4.3 | [Atp synthesis coupled electron transport](https://amigo.geneontology.org/amigo/term/GO:0042773) |
| 8.9E-06 | 18 | 44 | 4.3 | [Mitochondrial atp synthesis coupled electron transport](https://amigo.geneontology.org/amigo/term/GO:0042775) |
| 2.2E-05 | 17 | 43 | 4.1 | [Aerobic electron transport chain](https://amigo.geneontology.org/amigo/term/GO:0019646) |
| 9.8E-06 | 19 | 50 | 4 | [Oxidative phosphorylation](https://amigo.geneontology.org/amigo/term/GO:0006119) |
| 2.9E-05 | 18 | 49 | 3.8 | [Respiratory electron transport chain](https://amigo.geneontology.org/amigo/term/GO:0022904) |
| 3.9E-05 | 18 | 50 | 3.8 | [Electron transport chain](https://amigo.geneontology.org/amigo/term/GO:0022900) |
| 2.9E-05 | 21 | 64 | 3.4 | [Mitochondrial membrane organization](https://amigo.geneontology.org/amigo/term/GO:0007006) |
| 1.6E-07 | 39 | 140 | 2.9 | [Mitochondrial translation](https://amigo.geneontology.org/amigo/term/GO:0032543) |
| 2.1E-12 | 63 | 230 | 2.9 | [Mitochondrion organization](https://amigo.geneontology.org/amigo/term/GO:0007005) |
| 5.7E-05 | 31 | 126 | 2.6 | [Cellular respiration](https://amigo.geneontology.org/amigo/term/GO:0045333) |
| 2.2E-05 | 39 | 171 | 2.4 | [Mitochondrial gene expression](https://amigo.geneontology.org/amigo/term/GO:0140053) |
| 9.2E-06  Supplementary Table S4: list of downregulated genes | 103 | 646 | 1.7 | [Protein-containing complex assembly](https://amigo.geneontology.org/amigo/term/GO:0065003) |

| **Gene_Name_GFF** | **log2FoldChange** | **Pvalue** | **Protein Names** | **Function** |
| --- | --- | --- | --- | --- |
| **ZPS1** | -4.99808927 | 1.00877E-28 | Protein ZPS1 | Putative zinc-binding cell-wall/vacuolar protein; exact function unknown |
| **SPG1** | -3.619288043 | 5.45633E-07 | Stationary phase gene 1 protein | Stationary-phase associated protein; exact role unknown |
| **GPM2** | -1.561548949 | 7.52752E-07 | Phosphoglycerate mutase 2 | Putative phosphoglycerate mutase-like protein; function not well defined |
| **YBR285W (HAB1)** | -2.491256004 | 7.64701E-07 | YBR285W isoform 1 | Autophagy-related protein; binds ribosome; involved in autophagosome formation |
| **ASP3-1** | -2.915763568 | 1.15081E-06 | L-asparaginase II | Hydrolyzes L-asparagine → L-aspartate + NH₃; induced under nitrogen starvation |
| **CYC7** | -3.242196569 | 1.56532E-06 | Cytochrome c isoform 2 | Electron carrier in mitochondrial electron transport chain |
| **YGR174W-A** | -3.309987915 | 3.99782E-06 | Iso-2 cytochrome c | Likely a cytochrome c–related protein; function not well defined |
| **SPG4** | -3.397099725 | 5.92546E-06 | Stationary phase protein 4 | Stationary-phase induced protein; function unknown |
| **EGO4** | -3.505856702 | 6.55662E-06 | Exit from rapamycin-induced growth arrest protein | Member of EGO/GSE complex; regulates microautophagy and TORC1 signaling |
| **SUE1** | -2.242726368 | 1.58347E-05 | Protein SUE1, mitochondrial | Mitochondrial protein involved in protein quality control / protein degradation |
| **UIP4** | -1.573174773 | 1.75392E-05 | ULP1-interacting protein 4 | Unknown; interacts with SUMO protease ULP1; likely involved in SUMO pathway |
| **YAP6** | -2.502987592 | 1.87499E-05 | Transcription factor YAP6 | Transcriptional repressor involved in osmotic & stress-related gene regulation |
